# Human-specific progenitor sub-domain contributes to extended neurogenesis and increased motor neuron production

**DOI:** 10.1101/2022.07.05.498885

**Authors:** Sumin Jang, Elias Gunmit, Hynek Wichterle

**Author notes:** Corresponding Authors, Hynek Wichterle, Sumin Jang.

## Abstract

Neurogenesis lasts ~10 times longer in developing humans compared to mice, resulting in >1000-fold more neurons in the human central nervous system. Expansion of human neocortex has been in part attributed to the population of outer radial glia and amplifying progenitors that increase the output of neurogenic lineages. However, outer radial glia appear to be absent in many regions of the developing human nervous system, prompting us to search for alternative populations of progenitors that contribute to the expansion of human neurogenesis in one such region - the developing spinal cord. To this end, we performed high-temporal resolution single-cell expression analysis of human and mouse motor neuron progenitors generated from pluripotent stem cells in vitro. Alignment of human and mouse data using canonical correlation analysis identified “human-specific” progenitor clusters characterized by early co-expression of NKX2-2 and OLIG2 that lacked an orthologous murine counterpart. A matching progenitor population has been previously described in the human embryonic spinal cord^1^, but its function remained unknown. Our lineage tracing analysis demonstrates that these cells function as ventral motor neuron progenitors (vpMNs), but in contrast to classical pMNs, vpMNs exhibit increased Notch activity and generate motor neurons in a delayed and protracted manner. Concomitantly, vpMNs undergo more rounds of cell division before undergoing neurogenesis, leading to ~2-fold increase in total motor neuron output, and contributing preferentially to later-born, limb-innervating motor neuron subtypes. Thus, instead of relying on transit-amplifying progenitors, human spinal cord evolved a novel progenitor subdomain that extends timescales and expands output of human motor neurogenesis.

## INTRODUCTION

Vertebrate organisms exhibit remarkable diversity in the size and complexity of their central nervous systems (CNS). These size differences are reflected in the wide ranges in the duration of neurogenesis. The human brain contains ~1000 times more neurons compared to the mouse brain, and the neurogenic period required to support its development is ~10 times longer ^2–4^. This indicates that neural progenitors and modes of their division have to differ in fundamental ways across vertebrate species. What underlies these remarkable differences in neurogenic timescale and neuronal output across species is a matter of intense investigation^5–7^.

Within the CNS, the neocortex has undergone the greatest degree of expansion in human evolution^4,5,8,9^. Investigation of the primate neocortex revealed that unlike the rodent brain, primate ventricular zone progenitors give rise to expanded populations of outer radial glia (ORG) and intermediate progenitors that amplify neuronal output^10–12^. Whether similar, amplifying progenitors exist in other parts of the nervous system is not known. To address this question, we sought to investigate mechanisms underlying increased neurogenic output during human spinal cord development and motor neuron production.

Spinal motor neurons (MN) innervate muscles and sympathetic ganglia, controlling body posture, voluntary movement and the function of visceral organs ^13,14^. The human spinal cord generates >4 times the number of motor neurons compared to mouse **(Supplemental figure 1A)**, concordant with the increase in the size of our spinal cords and complexity of human motor skills^15–20^. Accordingly, human motor neuron genesis spans approximately 2 weeks from Carnegie stage 11 (post-conception day 26-30) to Carnegie stage 17 (post conception day 41-44), compared to ~2 days from embryonic day 9.5 to 11.5 in mouse^15,21^. Spinal motor neurogenesis from human and mouse pluripotent stem cells in vitro follows a similar timeline. In culture, mouse postmitotic neurons are born between day 5 and 7 in vitro, while the bulk of human motor neurogenesis occurs between days 10 to 20 of culture, and continues at lower rates well beyond.

Recently, it has been proposed that increased protein stability underlies the approximately 2.5-fold prolongation of key developmental processes associated with human versus mouse motor neuron genesis^15^. While this mechanism can explain the delay in the specification of human motor neuron progenitors, it is on its own insufficient to explain the ~7-fold increase in the motor neuron neurogenic period.

In order to identify the cellular and molecular factors underlying the expansion of human motor neurogenesis, we generated a high-temporal-resolution time-series of single-cell RNA-seq data of human and mouse cells undergoing in vitro motor neurogenesis. From this data, we identified orthologous as well as species-specific cell states. Among human-specific progenitors, we identified a population characterized by early co-expression of NKX2-2 and OLIG2, which has been reported in the developing human, but not mouse spinal cord^1^. Lineage tracing revealed that these cells give rise to motor neurons and therefore constitute a human-specific motor neuron progenitor population (vpMNs; ventral motor neuron progenitors). vpMNs exhibit increased Notch signaling activity and undergo delayed and protracted neurogenesis compared to the classical pMNs, ultimately expanding neuronal output by about two-fold. In summary, our findings establish that, unlike cortical neurogenesis where neuronal output is increased by outer radial glia and transit-amplifying cells, the human spinal cord evolved a distinct ventral progenitor sub-domain that contributes to the increased and prolonged production of spinal motor neurons.

## RESULTS

### Motor neuron differentiation involves several distinct progenitor cell states

To start delineating differences in rodent and human motor neurogenesis, we compared timelines of motor neuron production during in vitro differentiation of human and mouse pluripotent stem cells **(Supplemental figure 1B).** As reported previously, it took roughly 2-3 fold longer for motor neurons to first appear in human (day 11) compared to mouse (day 5) cell culture^15,19,20^ **(Figure 1A)**. Birthdating of spinal motor neurons by cumulative BrdU labeling at progressively later timepoints revealed that it takes ~40 hours for 70% of mouse motor neurons to be born, while it takes ~200 hours for 70% of human motor neurons to be born, corresponding to a 5-fold extension of the neurogenic period in human differentiation in vitro **(Figure 1B)**.

**Figure 1:**
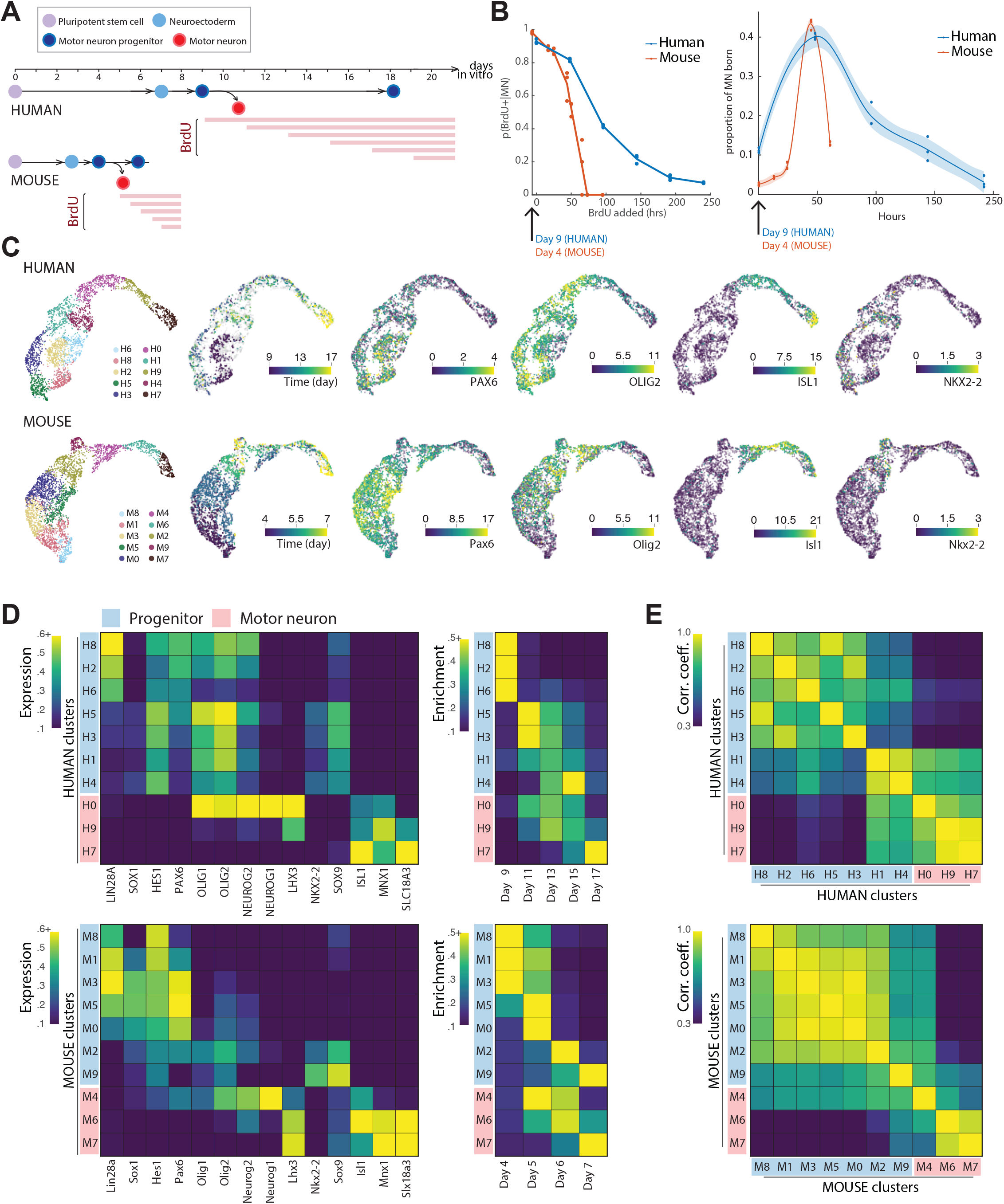
Temporal single-cell RNA-seq of in vitro mouse and human motor neurogenesis. **A)** Timeline of in vitro human and mouse motor neurogenesis and cumulative BrdU labeling. Red arrows denote timepoints selected for single-cell RNA-seq. **B)** Left: proportion of BrdU-positive MNs (y-axis) as a function of BrdU added at consecutive time points (x-axis) in cumulative BrdU labeling experiment. Right: birthcurve of human and mouse motor neurogenesis, as inferred from cumulative BrdU labeling experiment. **C)** UMAP (uniform manifold approximation and projection) plots of human and mouse single-cell gene expression profiles, colored by cluster identity, timestamp identity, and expression levels of key marker genes. **D)** Assignment of progenitor or motor neuron identity to cell clusters (rows) based on expression levels of marker genes (columns). **E)** Pearson correlation coefficients between expression profiles of high variance genes in human or mouse clusters.

Based on this timeline we selected days 4, 5, 6, 7 for mouse and days 9, 11, 13, 15, 17 for human differentiation, at which we examined cellular composition of cultures and the molecular identity of progenitors by single-cell RNA-seq. Human and mouse pluripotent stem cells were barcoded prior to differentiation using a lentiviral vector harboring constitutively expressed GFP with a 6bp barcode in its 3’ UTR region. Cell lines carrying distinct barcodes were differentiated in a temporally staggered manner to cover the selected timepoints, dissociated, pooled and processed on the 10x Genomics platform. In total, we obtained 3090 human and 3605 mouse high-quality single cell expression profiles across two biological replicates each, and assigned the barcoded timestamp to 33.5% and 60.9% of these, respectively **(Supplemental figure 1C, D)**.

Single cell expression profiles were processed and clustered using Seurat^22^,^23^, which yielded robust cell clustering configurations that remained >80% stable across a range of clustering parameters **(Figure 1C)**. Progenitor clusters were identified by high expression of dorso-ventral progenitor markers PAX6^24,25^ (low in motor neuron progenitors and increases dorsally) OLIG2^26,27^(motor neuron progenitor marker) and/or NKX2-2 (p3 marker, also co-expressed with OLIG2 in later-appearing oligodendrocyte precursor cells)^28^ and motor neuron clusters by high ISL1, MNX1 and/or SLC18A3 (VACHT) expression^29–31^ **(Figure 1D)**. For either species, we identified seven progenitor and three motor neuron clusters. Based on the timestamps, progenitor clusters could be further subdivided into early progenitors (present primarily in day 4 mouse and day 9 human cultures), mid-stage progenitors (day 5 mouse and day 11 human cultures) and two late-stage progenitors (day 6-7 mouse and day 13-15 human cultures). Early progenitor clusters (H8, H2, H6, M8, M1, M3) were enriched for LIN28A^32^ and PAX6, and mid-stage clusters (H5, H3, M5, M0) were enriched for OLIG2. While we did not identify clusters that correspond to the ventralmost p3 progenitors expressing NKX2-2 in the absence of OLIG2, we identified several progenitor clusters that co-express NKX2-2 and OLIG2. In mouse, this cluster (M2) is enriched in late cultures (day 6-7) and likely represent oligodendrocyte precursor cells (OPCs)^27,33,34^. In human, two out of the four NKX2-2/OLIG2 co-expressing clusters (H5, H3) were most enriched in mid-stage cultures on day 11 and the other two (H1, H4) are most enriched for later time-points: day 13 and 15, respectively **(Figure 1D)**.

In both human and mouse datasets, one postmitotic cluster (M0, H4) was identified as nascent motor neurons, co-expressing motor neuron progenitor (pMN) markers OLIG2 and NEUROG2 with postmitotic motor neuron (MN) markers ISL1 and MNX1. Two clusters were identified as postmitotic motor neurons in either species, characterized by high ISL1, MNX1 and SLC18A3 levels. Motor neuron clusters could also be temporally ordered according to differential timestamp enrichment: nascent motor neuron clusters H4 and M0 were enriched for day 11 and day 5-6, respectively. H9 and M6 were enriched for day 6 and days 13/15, respectively. Clusters H7 and M7 were enriched for the latest time point (day 7 and day 17, respectively). In the human dataset, postmitotic motor neuron clusters H9 and H7 are characterized by high expression of LHX3 and FOXP1 that, in the brachial spinal cord, specifically label medial motor column (MMC) and lateral motor column (LMC), respectively^30,35–37^. As LMC motor neurons are born after MMC motor neurons^30^, this indicates that H9 and H7 are enriched for different subtypes as well as different post-mitotic maturation states.

### Manifold alignment reveals orthologous and species-specific cell states

To compare the human and mouse cell clusters in an unbiased way, without having to rely on select marker genes or direct orthologous gene conversions between the two genomes, we used canonical correlation analysis (CCA)-mediated manifold alignment^22,23,38,39^. This analysis yielded a subspace of gene expression onto which both human and mouse cells can be mapped and clustered, ultimately providing a set of common clusters comprised of both mouse and human cells **(Figure 2A, B)**.

**Figure 2:**
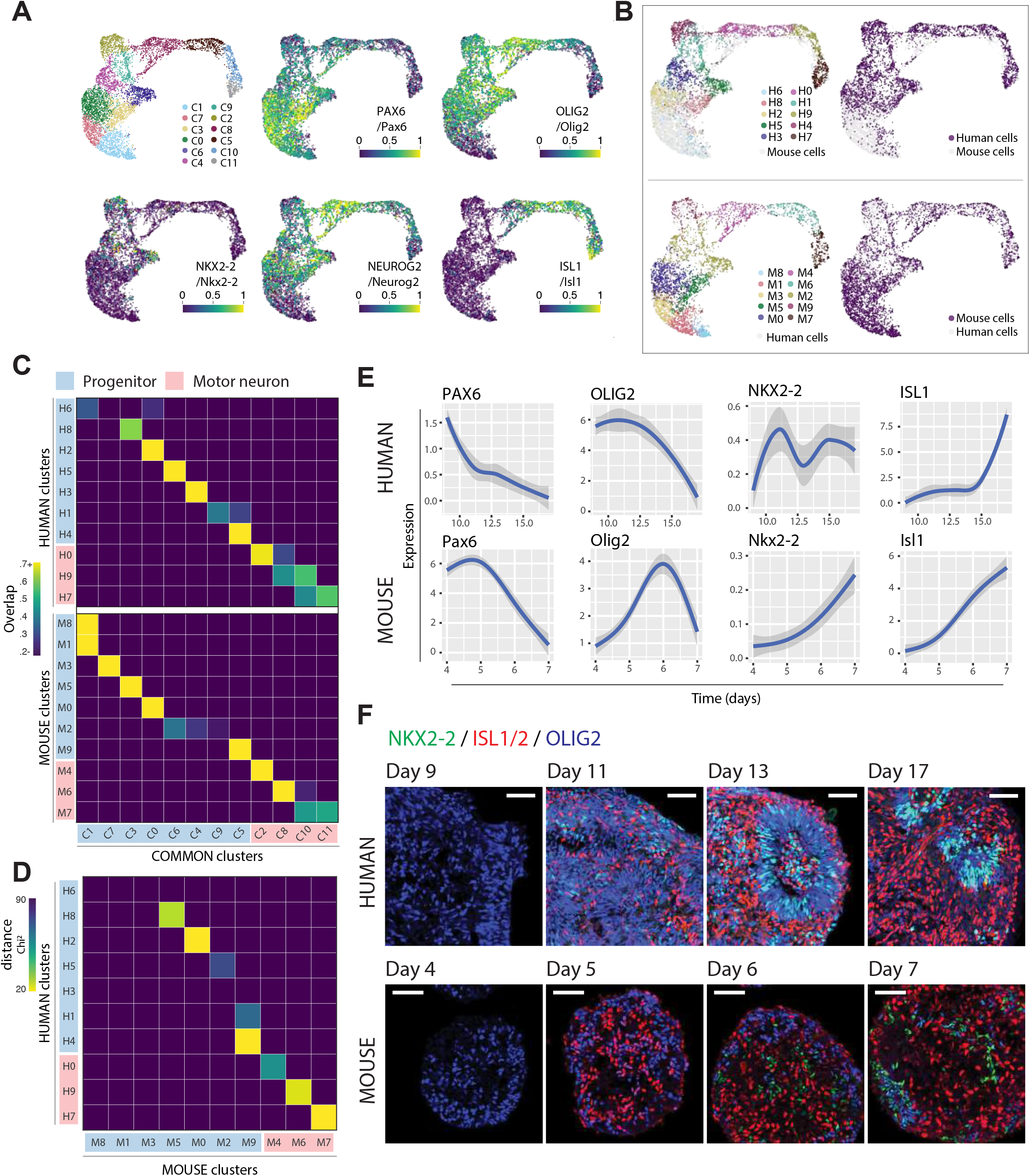
Alignment of human and mouse single-cell RNA-seq data reveals orthologous as well as species-specific cell states. **A)** UMAP plots of combined human and mouse single-cell expression profiles following canonical correlation analysis (CCA)-based integration. Cells are colored based on cluster identity or expression level of key marker genes. **B)** The same UMAP plots as in Figure 2A, with only human (top) or mouse (bottom) cells colored. **C)** Normalized overlap between individual human or mouse clusters (rows) to the set of common clusters (columns), as defined by clustering following integration. **D)** Chi-square distance between human and mouse clusters. Smaller distances indicate greater similarity in overlap distribution across common clusters **E)** Expression dynamics of key marker genes in human and mouse shows that human NKX2-2 expression is uniquely characterized by upregulation prior to peak neurogenesis (ISL1 upregulation). **F)** Immunofluorescence labeling of human and mouse embryoid bodies for NKX2-2, ISL1/2 and OLIG2 across different timepoints. Human NKX2-2 is first detected at the onset of neurogenesis (day 11) (Scale bars = 50um).

We found that cells assigned originally to progenitor and postmitotic motor neuron clusters distributed across distinct sets of common clusters **(Figure 2C)**. Human and mouse MN clusters (nascent, young and older) mapped onto the same set of four common clusters: the vast majority of nascent MNs from either species occupied one common cluster, whereas young and older MN cells tended to distribute across the remaining three common clusters. OPC-like late-stage human and mouse progenitor clusters (M9 and H4) also mapped faithfully onto the same common cluster. Using this logic, we quantified the distance between a given pair of human and mouse clusters by measure of how similarly the cells in either human or mouse cluster distributed across the set of common clusters **(Figure 2D)**. Orthologous cell states were defined as a human and mouse cluster pair with a mutually smallest distance (of chi^2^ < 60) to each other **(Figure 2D)**.

From this analysis, we found that about half of the early-to-mid stage progenitor clusters had clear orthologous counterparts, and the other half appeared to occupy largely human-dominated or mouse-dominated common clusters, indicating these cell states are “species-specific”. Mouse-specific (i.e., lacking an orthologous counterpart in the human dataset) clusters were early-stage progenitor clusters (M8, M1, M3), likely because the earliest time point collected for mouse (day 4) represents a less mature state compared to that of human cultures (day 9). Similarly, we identified four human-specific progenitor clusters – one early-stage (H6), two mid-stage (H3 and H5), and one late-stage (H1). H6 cluster is characterized by high expression of Pax6, a marker of neuroepithelial cells appearing in human neural differentiation earlier than in mouse^40^. The other human-specific clusters (H5, H3, H1) were characterized by co-expression of NKX2-2 and OLIG2. Of these, the mid-stage H3 cluster showed a markedly larger distance to any given mouse cluster **(Figure 2D)**.

The species-specific presence of early spinal progenitors co-expressing OLIG2 and NKX2-2 has been previously observed in human spinal cord in vivo and in differentiating stem cells in vitro^1,16^. We found that our human cultures exhibited similar characteristics. Whereas Nkx2-2 mRNA peaks after expression of motor neuron marker Isl1 in mouse cultures, human NKX2-2 expression shows much earlier upregulation that precedes OLIG2 downregulation **(Figure 2E)**. Immunocytochemical analysis of differentiating mouse and human stem cells confirmed the presence of human-specific OLIG2 and NKX2-2 double-positive progenitors in human cultures as early as on day 11 of differentiation, when ISL1-expressing motor neurons first appear **(Figure 2F)**.

To determine whether the H3 population corresponds to the species-specific, NKX2-2 and OLIG2 coexpressing progenitors in the early human embryonic spinal cord, we performed alignment of our data with existing scRNA-seq data from human Carnegie stage 12 spinal cord or mouse E9.5-E10.5 cervicothoracic spinal cord^16,41^ **(Materials and methods)**. The alignment revealed that the human-specific in vitro progenitors have matching human in vivo counterparts. In contrast, H3 cells lacked an orthologous counterpart in the in vivo mouse dataset **(Supplemental figure 2A)**. Interestingly, the human-specific progenitor clusters displayed poor overlap with NKX2-2-high, presumptive p3 progenitors in vivo and aligned more closely with an OLIG2-high, NKX2-2-low progenitor cluster (HF0), suggesting that these cells might belong to the MN lineage. Based on this analysis, we concluded that the identified in vitro human-specific progenitors exist in the developing human spinal cord in vivo, but not in mouse, and are transcriptionally distinct from pMNs and later appearing OPCs that are common to both mouse and human.

### Early NKX2-2/OLIG2 co-expressing cells are motor neuron progenitor cells

Differential gene expression analysis between human-specific and non-human-specific progenitor cells identified NKX2-2 as the best marker for the human-specific progenitor cells **(Figure 3A)**, exhibiting robust expression in human-specific progenitors and absence in other progenitor populations present in day 9-11 human cultures. To directly investigate the developmental fate of human specific progenitors, we engineered a human iPSC line with P2A-CRE-ERT2 inserted in-frame in front of the NKX2-2 stop codon. We also introduced a CAGGS-lox-STOP-lox-tdTomato cassette in the AAVS1 locus, to facilitate tamoxifen-inducible labeling of NKX2-2 expressing cells and their progeny **(Figure 3B)**. In order to selectively label early-mid stage NKX2-2-expressing human-specific progenitors, the differentiating cultures were pulsed with 200nM 4-hydroxytamoxifen (4OHT) between day 9 and 11 **(Supplemental figure 3A)**. This resulted in robust RFP expression detected as early as day 11, at which point the majority of labeled cells (>70%) also expressed NKX2-2 protein **(Figure 3C)**. By day 13, the number of RFP^+^ cells that were positively immunolabeled for NKX2-2 dropped below 60%, suggesting that the labeled progenitors started to differentiate into postmitotic NKX2-2 negative cells, and/or downregulated NKX2-2 expression (**Supplemental figure 3B**).

**Figure 3:**
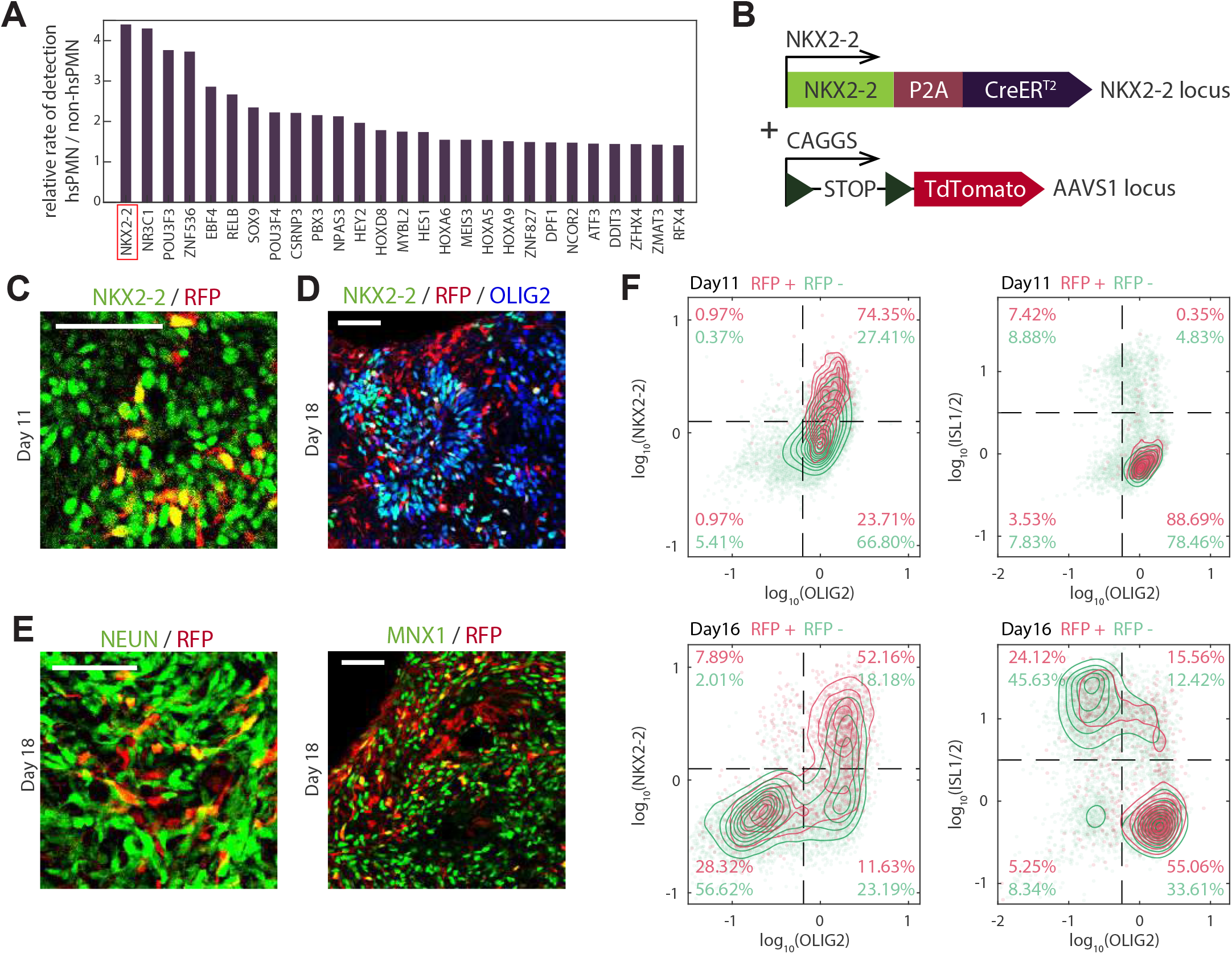
Generation of an NKX2-2-driven Cre-ERT2 human iPS cell line. **A)** The relative rates of detection (y-axis) in human-specific progenitors (vpMN) versus pMN cells shows that NKX2-2 expression is highly selective to human-specific progenitors. **B)** Schematic of NKX2-2-P2A-CreERT2 human iPS cell line design. **C)** Day 11 human cells immediately after 48 hours of 4-OHT treatment immunostained for NKX2-2 shows that the majority of RFP-positive cells are NKX2-2-positive. (Scale bar = 50um) **D,E)** Day 18 human cells after 4OHT treatment immunostained for pan-neuronal marker NEUN, NKX2-2, OLIG2 and MNX1. NEUN^+^/RFP^+^ and MNX1^+^/RFP^+^ cells indicate that human-specific progenitors give rise to motor neurons by day 18, although many RFP^+^ cells still remain progenitor-like (OLIG2^+^ and/or NKX2-2^+^). (Scale bars = 50um) **F)** Flow-cytometry analysis of day 11 and day 18 RFP^+^ and RFP^-^ cells immunostained for NKX2-2, OLIG2 and ISL1/2.

To determine the lineage potential of NKX2-2 progenitor cells, we examined the identity of RFP-expressing cells on days 16 and 18, following a 48-hour 4OHT pulse from day 9 to 11 **(Figure 3D, E)**. Immunostained sections of day 18 cultures revealed that the majority (~60%) of RFP^+^ cells had differentiated into postmitotic neurons expressing pan neuronal marker NEUN **(Figure 3E)**. Importantly, most of NEUN-positive cells also expressed motor neuron markers ISL1 and/or MNX1 **(Supplemental figure 3C, D)**, indicating that human-specific, NKX2-2-expressing progenitors are bona-fide motor neuron progenitors. To rigorously quantify the proportion of RFP-positive cells giving rise to motor neurons, we dissociated day 16 cultures and quantified cells immunostained for NKX2-2, OLIG2, and ISL1/2 by flow cytometry **(Figure 3F)**. This revealed that at day 16, ~60% of RFP-positive cells were still NKX2-2-positive, while the remaining ~40% differentiated into ISL1/2^+^ motor neurons.

Henceforth we refer to the human-specific, early OLIG2 and NKX2-2 double-positive cells as ventral motor neuron progenitors (vpMNs), based on the location of these progenitors in the ventral aspect of the OLIG2-positive motor neuron progenitor (pMN) domain in the human embryonic spinal cord^1^.

### vpMN cells exhibit delayed neurogenesis and increased motor neuron output

In order to investigate in more detail how vpMN cells differ from classical OLIG2^+^/NKX2-2^-^ progenitors (pMNs), we performed differential gene expression analysis between the two populations **(Materials and methods)**. Strikingly, we found that Notch signaling-related genes were some of the most differentially expressed between vpMN and pMN cells **(Figure 4A)**. Whereas genes that indicated lower Notch activity (NEUROG1, NEUROG2 and DLL1 were downregulated in vpMN cells, genes like NOTCH1, along with downstream effectors HES1/4, HEY1/2 were upregulated **(Figure 4B)**, indicating the vpMN cells had higher Notch signaling activity compared to pMN cells. Because Notch signaling has been shown to promote neural progenitor identity and inhibit neurogenesis^42^,^43^, we hypothesized that vpMN cells may be biased towards self-renewal and undergo delayed neurogenesis compared to pMN cells.

**Figure 4:**
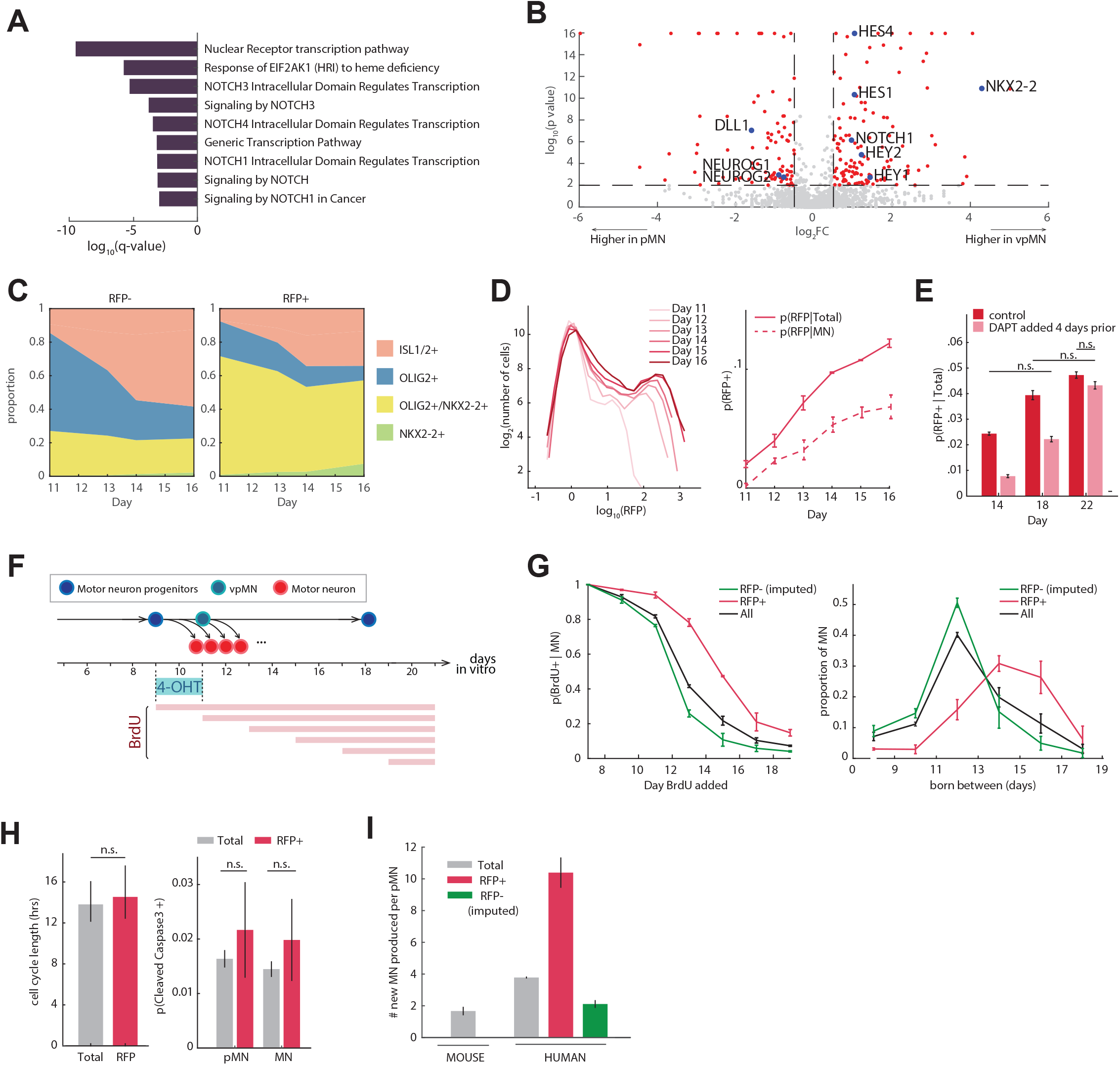
vpMN cells undergo delayed and protracted motor neurogenesis. **A)** Top enriched pathways in Reactome analysis of genes upregulated in vpMN (p-value <= 0.0001). **B)** Differential gene expression analysis between vpMN (H3) and pMN (H2 and H8) cells. **C)** Changes in the proportion of pMN (OLIG2^+^, ISL1/2^-^, NKX2-2^-^), MN (ISL1^+^), OPC or vpMN (NKX2-2^+^, OLIG2^+^, ISL1/2^-^), and NKX2-2^+^ cells over time in RFP-negative (left) and -positive (right) populations. RFP^+^ cells show relative depletion of MNs, especially during early neurogenesis (day 11-12). **D)** Left: Log-scale histogram of RFP expression over time shows that the proportion of RFP^+^ cells increases over time. Right: The proportion of RFP^+^ cells (y-axis) in both total and MN populations increases in a monotonic manner. **E)** RFP^+^ proportions stabilize upon DAPT addition to drive cells out of cell cycle (n.s. = non-significant; all pair-wise measurements are significantly different p <= 0.05 unless otherwise stated). **F)** Schematic of cumulative BrdU labeling assay to derive MN birthcurve. **G)** Left: Changes in the proportion of BrdU^+^ cells within MNs over time for RFP^+^, total and RFP^-^ (imputed) populations show a right-shift in the RFP^+^ BrdU curve, indicating that RFP^+^ remain mitotic until later stages on average. Right: Birthcurve of motor neurons in RFP^+^, total and RFP^-^ (imputed) populations shows that peak neurogenesis of RFP^-^ and RFP^+^ cells is between days 11-12 and days 13-17, respectively. **H)** Left: Estimation of cell cycle length based on EdU pulse-labeling shows that RFP^+^/OLIG2^+^/ISL1/2^-^ have similar cell cycle length compared to all OLIG2^+^/ISL1/2^-^ cells. Right: RFP^+^ and RFP^-^ cells display similar proportions of cleaved Caspase-3-positive progenitors and motor neurons. **I)** Average number of motor neurons produced per mouse or human (vpMN, pMN) progenitor cell.

To address this hypothesis, we first tracked how vpMN cells differentiate over time. We collected and dissociated embryoid bodies every 24 hours following 4OHT removal on day 11, stained cells for ISL1, OLIG2 and NKX2-2, then assessed proportions of pMNs (OLIG2^+^, NKX2-2^-^, ISL1^-^), motor neurons (ISL1^+^), and NKX2-2^+^/OLIG2^+^ cells (comprised of vpMN and later-appearing OPCs) over time in RFP^+^ and RFP^-^ populations via flow cytometry **(Figure 4C, Supplemental figure 4A)**. As expected, the proportion of progenitors progressively decreased, with a concomitant increase in the number of postmitotic motor neurons in both RFP^+^ and RFP^-^ populations **(Figure 4C)**. However, the proportion of progenitors and neurons differed between the two cell populations, with a larger fraction of RFP^+^ cells retaining progenitor identity throughout all time points, but especially so during the early stages of neurogenesis (day 11-13). This raised the possibility that RFP^+^ progenitors have an increased propensity to remain mitotic at the expense of undergoing neurogenesis. If this were the case, one would predict that the overall proportion of RFP^+^ cells would increase in the cultures over time. Indeed, we observed a monotonic increase in the proportion of RFP^+^ cells in the total population, as well as within total and newborn (born within the last 48 hours, as determined via BrdU labeling) MNs over time **(Figure 4D, Supplemental figure 4C)**. In contrast, when cells were treated with gamma secretase-inhibitor DAPT, which blocks Notch signaling and drives progenitor cells toward neurogenesis, the proportion of RFP^+^ cells no longer increased, but remained stable **(Figure 4E, Supplemental figure 4B)**.

To further evaluate the kinetics of motor neurogenesis in the vpMNs, we performed cumulative labeling of proliferating cells by adding BrdU to cultured cells on days 9, 11, 13, 15, 17, or 19, and terminated all samples on day 21 **(Figure 4F, Supplemental figure 4D)**. Based on this labeling strategy we calculated the proportion of day 21 MNs that became postmitotic prior to the addition of BrdU. This revealed a striking difference in the motor neuron birth curve of vpMN cells: whereas the overall motor neurogenesis peaked between days 11-13 and sharply decreased thereafter, neurogenesis in RFP^+^ cells steadily increased until day 13-15, remained high at day 17, and dropped only by day 19 **(Figure 4G)**. Considering that RFP^-^ cells contain a mixture of vpMN cells that failed to undergo Cre-dependent recombination as well as true pMN cells (successful recombination occurred in only 3-5% of NKX2-2 expressing cells), we imputed the birthcurve of true pMN cells (imputed RFP^-^ cells), based on the observation that 20-30% of OLIG2-expressing progenitors also express NKX2-2 **(Materials and methods)**. This revealed an even starker contrast in the birth curves of the two lineages. Whereas only 25% of pMNs were mitotic by day 13, >78% of vpMN cells remained mitotic on the same day **(Figure 4G)**.

Together, these observations indicate that vpMNs exhibit significantly delayed and protracted kinetics of neurogenesis. Considering that we did not observe significant differences in cell death or cell division rate among RFP^+^ and RFP^-^ cells **(Figure 4H)**, we reasoned that this delay in neurogenesis could allow vpMNs to undergo additional rounds of cell division and effectively amplify their neurogenic output (total number of neurons produced).

To directly compare the neurogenic output of vpMN and pMN cells in vitro, we seeded uniformly-sized embryoid bodies in round-bottom wells and counted the average number of RFP^+^ or RFP^-^ progenitors (OLIG2^+^, ISL1/2^-^) per embryoid body at day 11 (beginning of neurogenesis), as well as the average number of RFP^+^ and RFP^-^ motor neurons (MNX1^+^ and/or ISL1^+^) at day 18 (tail-end of neurogenesis). We also performed the same experiment with mouse cells, counting progenitors on day 5 and motor neurons on day 7.5 **(Supplemental figure 4E)**. This analysis revealed that while mouse motor neuron progenitors give rise on average to ~1.6 motor neurons, human progenitors give rise on average to ~4 motor neurons (**Figure 4I)**. Within human progenitors, vpMN cells produced on average >10 motor neurons, compared to an average of ~2 motor neurons produced by pMN cells **(Figure 4I)**. This difference in neurogenic output between the two lineages means that, despite their relatively low number compared to pMNs, vpMNs generate ~50% of all motor neurons observed in day 18 embryoid bodies.

### vpMN cells preferentially contribute to late-born, limb-innervating motor neurons

In vitro-differentiated human pluripotent stems cell yield diverse motor neuron subtypes, including LHX3-expressing axial MMC (medial motor column) motor neurons and FOXP1-expressing limb innervating LMC (lateral motor column) motor neurons, and LHX3/FOXP1-negative HMC (hypaxial motor column) motor neurons **(Figure 5A, Supplemental figure 5A)**^20,44,45^

**Figure 5:**
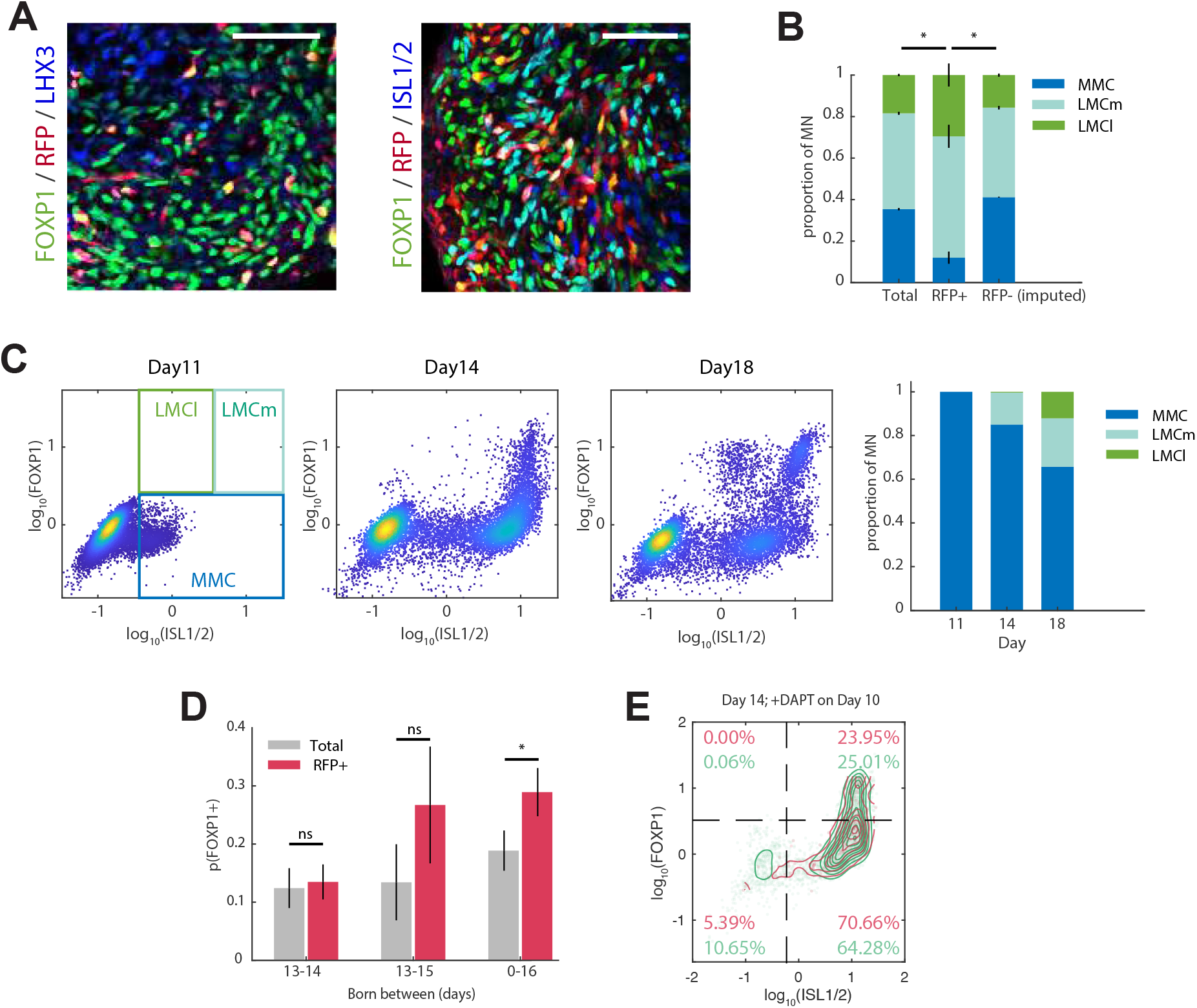
vpMN cells are biased toward later-born, limb-innervating motor neuron subtype fates due to delayed neurogenesis. **A)** Day 18 human cultures immunostained for LHX3, FOXP1 and ISL1/2 show that RFP+ cells are found in both MMC (LHX3^+^) as well as LMC (FOXP1^+^) cells. (Scale bars = 50um) **B)** The relative proportions of MMC, LMCm and LMCl subtypes in day 20 RFP^+^, total and RFP^-^ (imputed) populations following late DAPT treatment on day 15 shows that RFP+ cells are enriched for LMC subtypes. **C)** Flow cytometry analysis of cells immunostained for ISL1/2 and FOXP1 shows temporal specification of MMC, LMCm and LMCl subtypes. **D)** The proportion of FOXP1^+^ cells among those that are born in a 24-hour, 48hour, or 16-day window, based on EdU and BrdU dual-labeling. **E)** Day 14 human cells treated with DAPT on day 10 show similar FOXP1 expression in RFP^+^ and RFP^-^ populations.

To determine whether MNs derived from vpMNs contribute differentially to motor neuron subtypes, we stained cells for FOXP1 and ISL1 on day 17 following 4OHT treatment on day 9-11, and assessed the relative proportions of MMC/HMC (ISL1^+^/FOXP1^-^), LMCm (ISL1^+^/FOXP1^+^), LMCl (ISL1^low^/FOXP1^+^) in RFP-positive and -negative populations. Interestingly, we found that RFP^+^ vpMN-derived MNs displayed a significantly higher proportion of FOXP1^+^ cells than RFP^-^ cells **(Figure 5A, B)**.

Considering that ISL1^+^/FOXP1^+^ LMCm motor neurons and ISL1^-^/FOXP1^+^ LMCl motor neurons respectively appear in vitro around days 14 and 18, when vpMNs are neurogenic **(Figure 5C)**, we wondered whether vpMN-lineage motor neurons were enriched for later-born LMC subtypes due to their delayed timing of neurogenesis, or because vpMNs are intrinsically biased toward LMC fates. To test this, we pulsed 4OHT-treated (day 9-11) cells with EdU on day 13, followed by BrdU addition on day 14 to differentially label neurons born before, within and after the 24-hour window between days 13 and 14. As expected, the total percentage of FOXP1^+^ cells was significantly greater in the RFP^+^ population compared to background (29% in RFP^+^, 19% in total, p = 0.032). However, RFP^+^ and background cells that underwent neurogenesis in this 24-hour window (i.e., BrdU^+^, EdU^-^) had the same proportion of FOXP1^+^ cells (~12%). When this window was increased to 48 hours (BrdU added on day 15), FOXP1^+^ cells became more prevalent in the RFP^+^ population, whereas they remained at similar levels in the background population (25% in RFP^+^ and 12% in total, p = 0.127) **(Figure 5D)**. This suggests that vpMN and pMN cells undergoing neurogenesis at the same time have a similar probability of giving rise to FOXP1^+^ motor neurons.

To further test whether vpMN are intrinsically more biased toward LMC fates compared to pMN cells, we forced progenitors to prematurely differentiate into motor neurons by Notch inhibition with DAPT (gamma secretase inhibitor) on day 10. We found that both lineages simultaneously underwent early neurogenesis in response to DAPT, and became ~90% ISL1/2 and MNX1-positive within 96 hours. Further, when progenitors underwent premature and simultaneous neurogenesis in response to DAPT, both RFP^+^ and RFP^-^ cells harbored similar proportions of FOXP1^+^ cells **(Figure 5E)**, confirming our findings that both lineages give rise to similar proportions of LMC subtypes when normalized for timing of neurogenesis. Together, these results show that vpMN-derived motor neurons are enriched for FOXP1-expressing, limb-innervating LMC motor neurons as a direct result of their delayed neurogenic timeline, and not due to an intrinsic bias toward LMC fates.

## DISCUSSION

A key question in biology is why different organisms develop, reproduce and age according to different timescales. The biological “clock” for humans runs at a slower pace than that for mice. Many developmental processes (e.g., segmentation clock, cell cycle, differentiation, etc.) are protracted by roughly the same degree (~2.5-fold) between human and mouse, indicative of a general mechanism underlying the timing differences. Indeed, recent studies have shown that a slower protein turnover and/or metabolic rates in human may drive the 2.5-fold delay in the developmental timeline^15,46–48^.

However, not all developmental processes scale by this factor, as exemplified by neurogenesis that is protracted by a significantly larger factor. Specifically, cortical development that takes ~7 days in mouse (E11-E18) is >9 times longer, taking ~65 days in human (E40-E105)^49^. Similarly, spinal motor neurons are generated over a ~2-day period in brachial spinal cord spanning from day ~9 to day ~11 in mouse, while human motor neuron genesis extends from CS11 (day 27) to CS17 (day 41)^15^, a 7-fold increase in time.

In the neocortex, several key mechanisms were identified that contribute to the prolongation of the neurogenic period and increased neuronal output. The human fetal neocortex is characterized by an expansion of outer radial glial cells and amplifying progenitors that increase production of late born Layer 2/3 neurons^4,10,50^. In addition, human-specific paralogs of the NOTCH2 receptor gene (NOTCH2NL) increase Notch activity in neocortical progenitor cells, contributing to their expansion^51,52^, and increased activity in SHH, WNT and FGF leads to similar effects as well^53–56^. However, mechanisms controlling expansion of neurogenesis in other CNS regions is less well understood.

To identify mechanisms contributing to the prolongation of the neurogenic period in human spinal cord, we performed a detailed mapping of cellular diversity and molecular identity of human pluripotent stem cells differentiating to motor neurons in vitro. We identified transcriptionally distinct, human-specific subsets of neural progenitors that co-express NKX2-2 and OLIG2 and appear around the onset of motor neurogenesis. Such NKX2-2 and OLIG2 co-expressing cells were previously identified by histological analysis in the ventral aspect of the pMN domain of the spinal cord and in human single-cell RNA-seq data from Carnegie stage 12 embryonic samples^52^, whereas a similar cell cluster is absent in mouse embryonic data **(Supplemental figure 2A)**. However, it remained unknown whether these human specific cells are early specified oligodendrocyte progenitors (OPCs), motor neuron progenitors, or V3 interneuron progenitors. By integrating human and mouse single cell embryonic data, we found that the overall gene expression profile of vpMN cells showed much stronger correlation to that of other, NKX2-2-negative, pMN cells compared to that of later-stage, NKX2-2 and OLIG2 co-expressing OPC-like cell clusters **(Figure 1E)**.

To directly examine the fate of vpMNs we performed genetic lineage tracing, demonstrating that these cells are indeed motor neuron progenitors, expanding the timescales and output of human motor neurogenesis. We found that vpMN cells exhibit increased Notch signaling activity and undergo delayed and protracted neurogenesis compared to pMN cells. Because of these dynamics, the vpMN population is able to undergo further amplification during differentiation, with each vpMN progenitor giving rise to ~5 times as many motor neurons compared to pMNs. At the same time, these progenitors are biased toward generating later-born, limb-innervating LMC motor neuron subtypes. vpMN cells are therefore functionally analogous to outer radial glia (ORG) of the human neocortex, that also undergo expansion and delayed neurogenesis, thereby increasing the production of later-born Layer 2/3 pyramidal neurons. A key distinction between vpMNs and ORG cells is that vpMN cells are not descendants of the pMN population; instead, they constitute a distinct motor neuron sub-lineage that is spatially separated from pMNs.

It is of interest that within the lineage traced progenitor population (i.e., cells that expressed NKX2-2 at the time of tamoxifen administration) we observed a significant fraction of cells that expressed only OLIG2. This raises the possibility that double positive vpMNs downregulate NKX2-2 before motor neuron differentiation. Indeed, analysis of single cell RNA-seq data and immunocytochemical analysis revealed that nascent motor neurons co-expressing OLIG2 and ISL1 are present across all neurogenic timepoints **(Figure 1D, Supplemental figure 4A)**. In contrast, NKX2-2 and ISL1 co-expression is virtually absent (detected at <1% frequency) across all timepoints, even at late stages of motor neuron genesis when double positive vpMNs are giving rise to –late born motor neurons. This suggests that vpMN cells differentiating into motor neurons downregulate NKX2-2 expression prior to the activation of the postmitotic motor neuron gene expression program.

In addition to the OLIG2-only expressing progenitors, we identified a small population of NKX2-2-only expressing progenitors among the lineage traced cells in later stages of differentiation **(Figure 4C, Supplemental figure 4A)**. This raised the possibility that a subset of vpMN cells might downregulate OLIG2. Indeed, canonical correlation analysis of NKX2-2/OLIG2 double-positive cells present specifically in late-stage cultures mapped preferentially onto an NKX2-2-high, p3-like embryonic human cell cluster. Given these observations, NKX2-2 and OLIG2 co-expressing vpMN cells may represent a transient cell state that either downregulates NKX2-2 and adopts a “classical” pMN identity before differentiation into spinal motor neurons, or, albeit with a lower probability, downregulates OLIG2 and differentiates into V3 spinal interneurons.

Taken together, our studies identified a novel developmental strategy contributing to the extension and expansion of human embryonic spinal neurogenesis. We demonstrate that human-specific NKX2-2/OLIG2 double-positive cells appearing at early stages of neurogenesis are functionally and molecularly distinct from the later generated NKX2-2/OLIG2 double positive oligodendrocyte progenitors present in the mouse spinal cord. These cells serve as a reservoir of amplifying progenitors characterized by an increased activity of Notch signaling, biasing the cells to self-renewal (expansion) and delayed neurogenesis. The vpMN cells are positioned between the pMN and p3 domain in the developing spinal cord^1^, forming an evolutionarily novel progenitor domain. Whether these transcriptionally distinct progenitors generate fundamentally different and novel subtypes of motor neurons remains to be determined. We propose that the vpMN cells are molecularly transient, as they appear to lose NKX2-2 expression before their differentiation to motor neurons. This is consistent with the observation that vpMNs in the developing human spinal cord exhibit lower level of NKX2-2 compared to p3 progenitors^1,16^. Our analysis of human and mouse neurogenic output revealed that despite their smaller number, vpMN progenitors contribute significantly to the motor neuron numbers due to their increased self-renewal coupled with a delayed neurogenesis. Considering the evolutionary distance between mouse and human, it will be interesting to determine when during vertebrate evolution this population of progenitors emerged.

## MATERIAL AND METHODS

### Cell culture

#### Human cell culture

Human iPS cells (NCRM-1) were grown on Matrigel (Corning CLS354277)-coated plates in mTeSR-Plus (STEMCELL Technologies 100-0276), and differentiated according to previously established protocols^20^, without DAPT unless otherwise stated. For NKX2-2-CreERT2-dependent recombination, 200nM of 4-hydroxytamoxifen was added on day 9 and kept on for 48 hours. For cumulative BrdU assays, 10uM BrdU was added as early as day 9 and as late as day 19, kept on throughout the experiment and all harvested on day 21. For EdU/BrdU dual-labeling assays, 5uM EdU was added on day 13 for 24 hours, followed by 10uM BrdU which was kept on until the end of the experiment (day 16). DAPT was used at 10uM at various time points to drive progenitors out of the cell cycle.

#### Mouse cell culture

C57BL/6J Mouse ES cells were grown on gelatin-coated plates in LIF+2i^57^-containing basal media, and differentiated according to previously established protocols^19^. For cumulative BrdU assays, 10uM BrdU was added as early as day 4 and as late as day 7, kept on throughout the experiment and all harvested on day 8.5.

#### To generate uniformly-sized embryoid bodies

150 human or 200 mouse cells were seeded per ultra-low-adhesion round-bottom well (96-well format; Corning 7007) in day 0 differentiation media, and subsequent media changes were done individually for each well. In the case of human differentiations, embryoid bodies were transferred to larger vessels on day 16.

### Lentiviral barcoding and single-cell RNA-seq

Lentivirus was generated in HEK293T cells using a second-generation system with pMD2.G (Addgene 12259) and psPAX2 (Addgene 12260). Lentivirus harboring CAGGS-driven GFP with unique 6bp barcodes (“timestamps”) were used to transduce human iPS cells and mouse ES cells. 48-96 hours following transduction, GFP-positive cells were purified via FACS for continued culture and/or freeze-down. Each uniquely barcoded batch of cells was differentiated starting on different days, such that all timepoints could be harvested and processed for single-cell RNA-seq on the same day. Following differentiation, cells were dissociated, FACS-purified for GFP, pooled in equal parts across species and timepoints, and processed on a 10X Genomics V3 3’-end single-cell gene expression profiling platform at the Columbia University Sulzberger Genome Center. Single-cell RNA libraries were then sequenced on an Illumina Hi-seq 4000 at a depth of XX reads per cell.

### Processing and analysis of in vitro single-cell RNA-seq data

Single-cell RNA-seq reads were demultiplexed, aligned to hg38 (Genome Reference Consortium Human Build 38) and mm10 (Genome Reference Consortium Mouse Build 38) and quantified (barcode and UMI) using CellRanger version 3.1.0 (10X genomics). Prior to alignment, the reference fastq files were supplemented with GFP and barcode sequences for lentiviral timestamp detection and quantification. Following quantification, cell profiles with fewer than 1500 unique genes detected were discarded, along with cell profiles that indicated a fibroblast-like (COL1A2-high) or V2 interneuron lineage (high VSX2 or IRX3) identity. These cells were identified by preliminary clustering of each dataset individually, which made V2 or fibroblast-like cells, if present, easily distinguishable, as they would each be found in a distinct cluster. Remaining cell profiles were processed and clustered using Seurat v4.0.6: cell profiles were integrated from across three 10X runs, and cell cycle-related gene expression signatures were regressed out.

Integration of mouse and human datasets: a list of 1:1 orthologous genes was generated using the Ensembl biomaRt homolog dataset. This was then used to generate mouse and human sub-datasets of the original single-cell RNA-seq UMI counts dataset, where rows are all ortholog-matched. Ortholog-matched datasets were subsequently integrated and clustered together using Seurat v4.0.6.

### Processing and analysis of embryonic human and mouse single-cell RNA-seq data

Cell profiles (Human Carnegie stage 12^16^ and mouse E9.5 and 10.5^41^) were downloaded from EMBL’s European Bioinformatics Institute (mouse; E-MTAB-7320) and Gene Expression Omnibus (human; GSE171890) and processed using Seurat v4.0.6. Individual replicates and/or timepoints were integrated within each species and cell profiles were clustered following removal of cell cycle-related gene expression signatures. Following clustering, individual clusters that displayed high expression of OLIG2, NKX2-2, ISL1, or MNX1, but low expression of IRX3 and PHOXA2 were then selected and clustered again separately to generate final cluster assignments. Integration of in vivo and in vitro datasets were performed in the same manner as detailed above for integration of human and mouse datasets.

### Differential gene expression analysis

DEsingle^58^ v1.8.2 with standard parameters and default settings was used to generate a list of differentially expressed genes in pair-wise comparisons between two sets of single-cell gene expression profiles, and their p-values. False discovery rate-adjusted q-values were obtained using the Benjamini-Hochberg method with a threshold value of 0.05.

### CRISPR-mediated knock-in of P2A-CreERT2 downstream of NKX2-2

A single cut in the NKX2.2 3’UTR was made using transient transfection (Lonza Human Stem Cell Nucleofector Kit VPH-5012) of with CAGGS::Cas9-mCherry and gRNA expression vector^59^ (Addgene 41824) harboring guide RNA sequence: 5’-GGGGCCGCGAGTCTCGTTG-3’ (PAM: GGG), along with linearized donor vector harboring P2A-CreERT2 sandwiched between 800-bp homology arms for NKX2-2 exon 2 and 3’UTR regions. Following transfection, cells were recovered, re-plated at clonal density on mouse embryonic fibroblasts (irradiated; Gibco A34180), after which individual clonal colonies were picked and genotyped. Genotyping primers: 5’-CGAAAGCCCCCAAAAACCTG-3’, 5’-CGTAGAGTTCAGCCCTCTCC-3’

### Immunostaining

#### For flow cytometry or FACS

Embryoid bodies were dissociated using papain following the Neural Tissue Dissociation Kit – P (Miltenyi Biotec 130-092-628) and quenched with ovomucoid protease inhibitor (Worthington LK003182). Following dissociation, cells were washed twice with 1X PBS, fixed using 4% paraformaldehyde, washed with PBS, and then permeabilized with ice-cold methanol. Permeabilized cells were rehydrated with 1% BSA in PBS solution, and subsequently immunostained using a standard two-step immunofluorescence labeling method.

#### Sectioned embryoid bodies

Whole embryoid bodies were fixed in 4% paraformaldehyde, washed several times with PBS, then put through 30% sucrose prior to embedding in OCT and freezing. Frozen blocks were sectioned in a cryostat to 15um slices and mounted on microscope slides (Superfrost Plus, Fisher 12-550-15). Sections were rehydrated with staining solution (1% BSA, 0.1% Triton X-100 in PBS), then immunostained using a standard two-step immunofluorescence labeling method.

#### Antibodies used

ISL1/2 (Mouse, 1:100 DSHB 4D5 supernatant), ISL1/2 (Goat, 1:5000 Neuromics GT15051), MNX1 (Guinea pig, 1:100 from Jessell Lab), FOXP1 (Mouse, 1:400 Santa Cruz A-2 sc-398811), NKX2-2 (Mouse, 1:100 DSHB 74.5A5 supernatant), BrdU (Rat, 1:400 Abcam ab6326), LHX1/2 (Mouse, 1:100 DSHB 4F2 supernatant), NeuN (Rabbit, 1:1000 Millipore Sigma ABN78), OLIG2 (Guinea pig, 1:100 from Jessell Lab), Ki67 (Mouse, 1:100 BDPharmingen 550609), Cleaved Caspase3 (Rabbit, 1:200 Cell Signaling #9661), LHX3 (1:100 DSHB 67.4E12 supernatant)

#### EdU labeling

Click-iT Plus (Thermo Fisher C10640)

### Imaging

Images were acquired with 20X objective using confocal laser scanning microscope (LSM Zeiss 780).

### FACS and flow cytometry

BD FACSAria Cell Sorter equipped with 5 lasers (355nm, 405nm, 488nm, 561nm, 640nm) and 100um nozzle was used to acquire flow cytometry data (minimum 10,000 events per replicate) or FAC-sort cells.

### Estimation of cell cycle length based on EdU pulse

On day 14 of human culture, following 4-OHT pulse between days 9 and 11, 5uM EdU was added to the culture media for 5-23 hours, after which embryoid bodies were dissociated and fixed with paraformaldehyde (4%). Following fixation, cells were permeabilized with ice-cold 100% methanol, then rehydrated with 1% BSA in PBS and immunolabeled for OLIG2. EdU was then labeled using click chemistry. Using flow cytometry, the proportion of EdU-negative cells among OLIG2-positive cells, or *p*, was measured for both RFP^+^ and total populations. Assuming that cells undergo cell cycle in an asynchronous manner (i.e., each cell cycles independently of another), we calculated the average cell cycle length in a given population as:

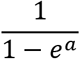

Where *α* is the slope given by log(*p*) = *α · t*, and *t* is the duration of EdU pulse.

### Reactome pathway enrichment analysis

Pathways enriched in differentially expressed genes were identified using the standard web version of Reactome analysis (reactome.org)^60^. Genes that were upregulated in vpMN relative to pMN with a q-value of less than 0.05 were used as input.

### Overlap between common and species clusters

The overlap between species cluster *i* and common cluster *j* was calculated as:

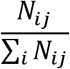

where *N_i,j_* is the number of cells with species cluster identity *i* and common cluster identity *j*.

Chi-square distance between human cluster *x* and mouse cluster *y* was calculated as:

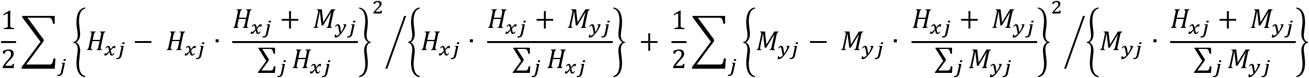

where *H_xj_* is the number of human cells with human cluster identity *x* and common cluster identity *j*, and *M_yj_* is the number of mouse cells with mouse cluster identity *y* and common cluster identity *j*.

### Imputing RFP-population characteristics

Because tamoxifen-induced recombination is low-efficiency (<1% of the population is RFP^+^ at day 11, even though 20-30% of cells are immunolabeled positively for NKX2-2), we imputed population characteristics using a conservative p(NKX2-2^+^) of 0.2. Given the proportion of a certain cell type (e.g., FOXP1^+^) in the RFP^+^ and total populations is *p* and *q*, respectively, its proportion in the RFP-negative population is:

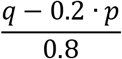

## AUTHOR CONTRIBUTIONS

S.J. and H.W. designed the study. S.J. performed experiments and data analysis. E.G. generated the human NKX2-2-CreERT2 iPS cell line. S.J and H.W. wrote the manuscript.

## ACKNOWLEDGEMENT

We would like to thank Dr. Grace Yeo and Dr. David Gifford for help with and access to their shared computing server, Erin Bush and Michael Finlayson of the Columbia University Sulzberger Genome Center for running the 10X single-cell library prep and sequencing, Mike Kissner and the CSCI Stem Cell Facility for assistance with flow cytometry. We would like to thank Sophia DiIorio for designing and testing sequencing primers for barcode analysis, and Dr. Gist Croft for immunostained human embryonic spinal cord images. We are also grateful to Dr. Carol Mason and Dr. Franck Polleux, the members of the Wichterle, Chaolin Zhang and Edmund Au labs for discussion and feedback. S.J. received support from the Columbia Stem Cell Initiative seed grant. H.W. is a recipient of the Jerry and Emily Spiegel endowed chair. This work was supported by NIH R01NS109217; R01NS116141; R01NS089676 and Project ALS.

## COMPETING INTERESTS

The authors declare no competing interests.

## SUPPLEMENTAL FIGURE LEGENDS

**Supplemental figure 1:**
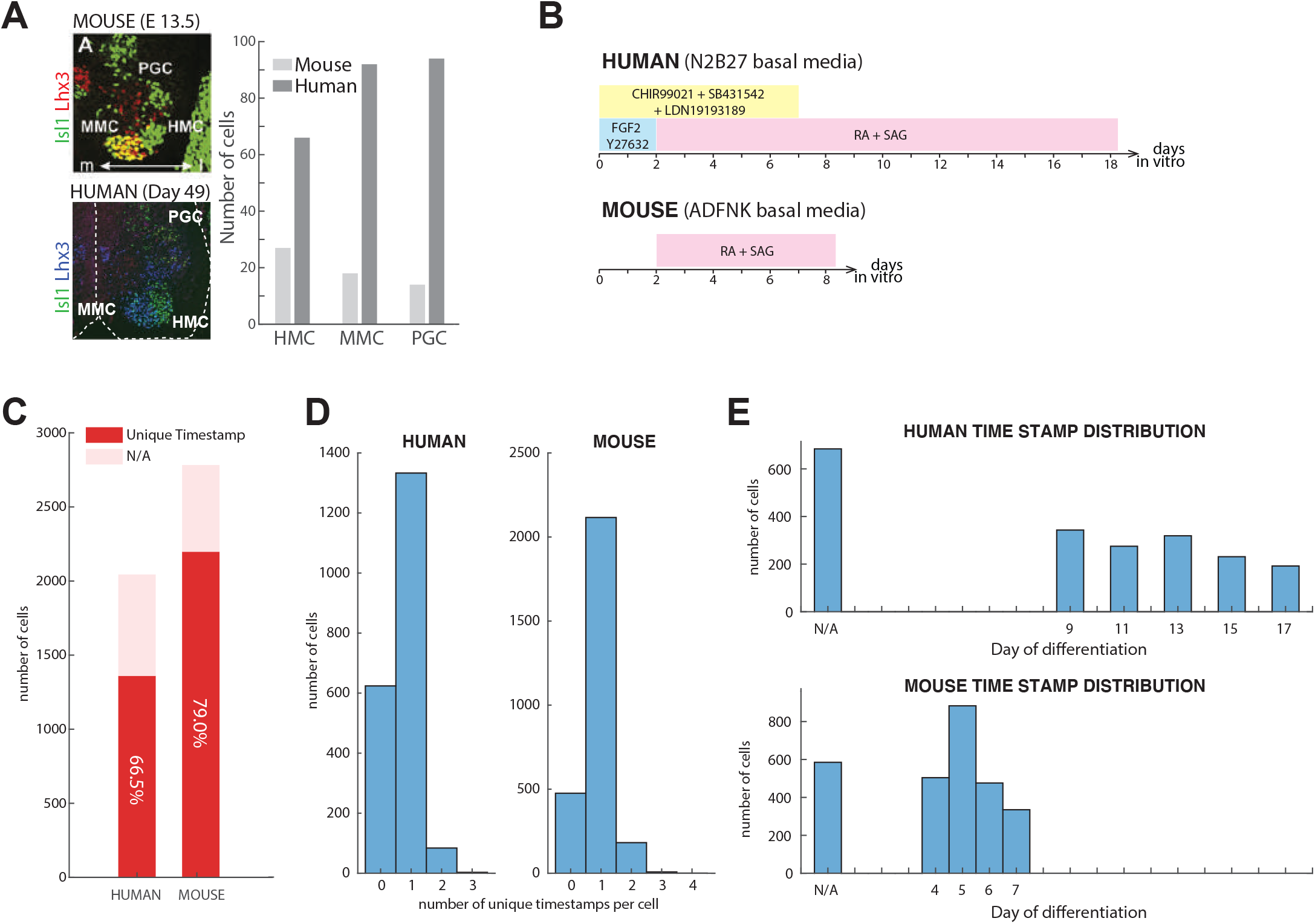
**A)** Thoracic sections of E13.5 mouse^61^ and day 49 human spinal cord shows increased numbers of all three motor neuron subtypes in human compared to mouse (medial motor column = MMC; hypaxial motor column = HMC; pre-ganglionic column = PGC). **B)** Culture conditions for motor neuron differentiation. **C)** Numbers of single-cell profiles (successfully time-stamped and not) from second batch of 10X single-cell gene expression profiling. Note, the first batch of human cells were not virally time-stamped, leading to overall lower proportion of time-stamped human cells. **D)** Distribution of the number of unique timestamps per cell shows that the majority of cells have one unique timestamp. **E)** Numbers of single-cell profiles obtained for each time-point, as determined by time-stamps.

**Supplemental figure 2:**
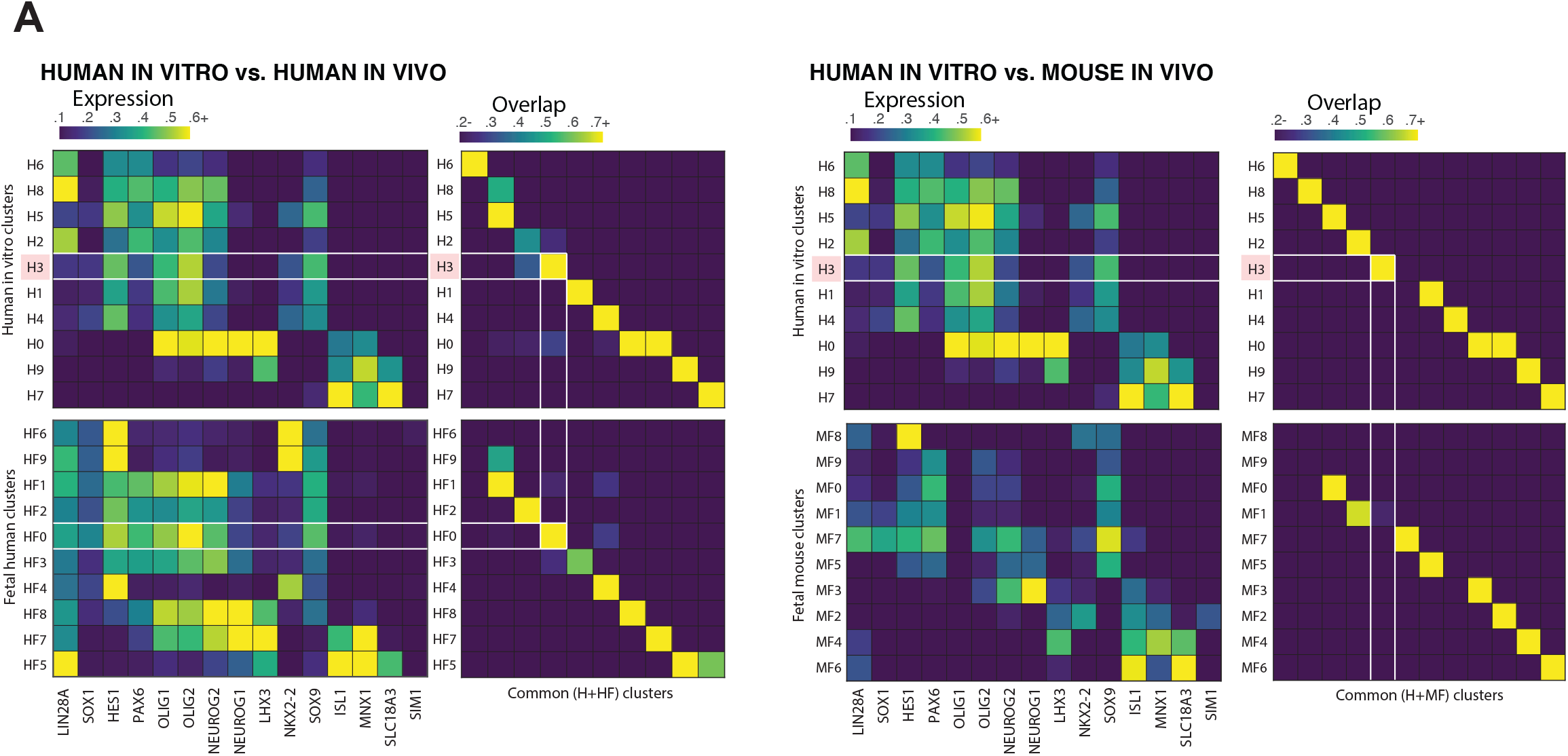
**A)** Left: Alignment of human in vitro (top) and human CS12 embryonic (bottom) single-cell RNA-seq samples shows that H3 cells map onto a distinct cluster found in vivo (cluster HF0). Right: Alignment of human in vitro (top) and mouse E9.5-10.5 spinal cord shows that H3 cells show poor overlap with all embryonic mouse clusters, suggesting that H3-like cells are found in human (but not mouse) embryonic spinal cords.

**Supplemental figure 3:**
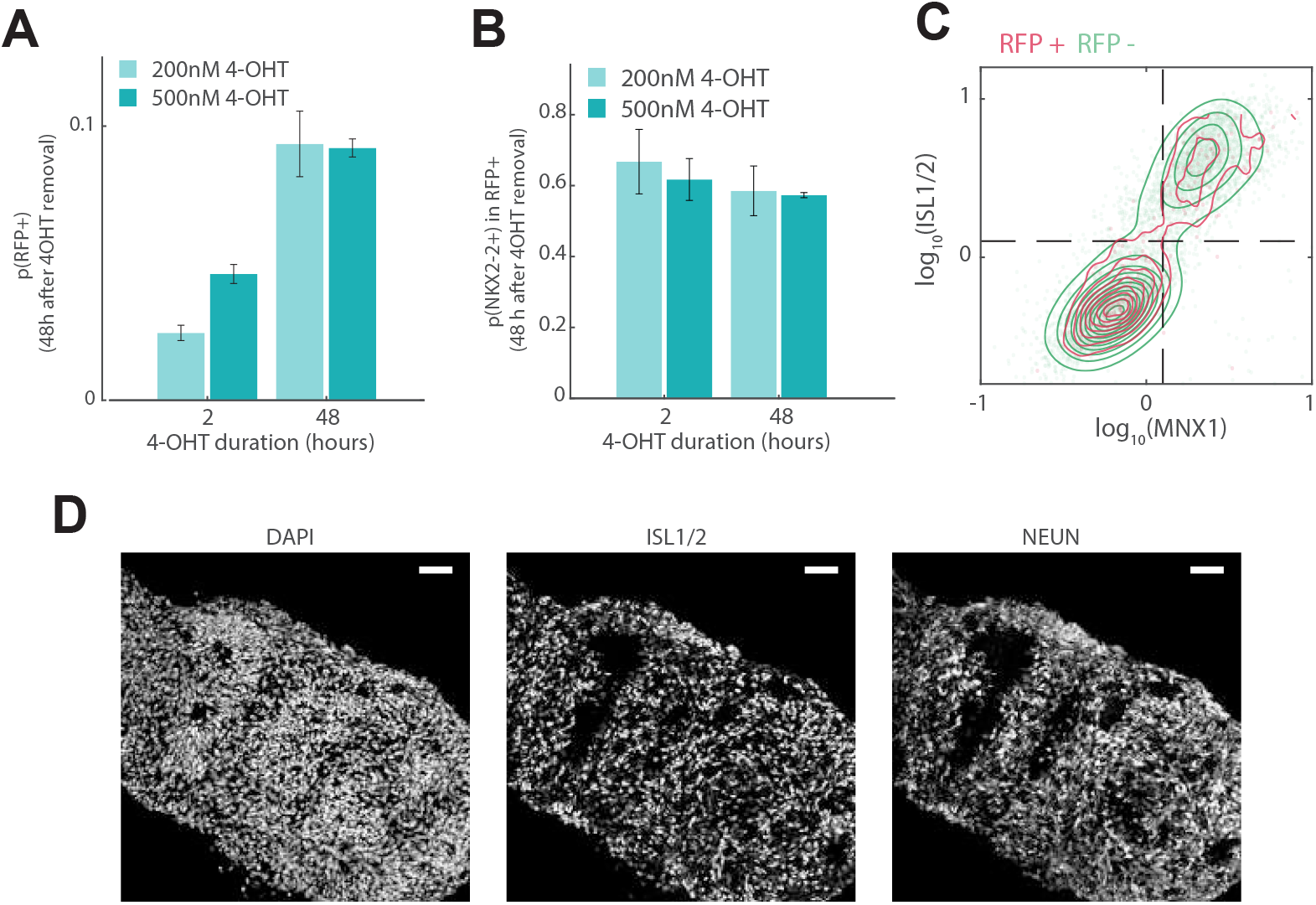
**A)** Varying lengths and concentrations of 4-OHT pulse affect the efficiency of recombination-based RFP expression. **B)** Proportions of NKX2-2-positive cells within RFP-positive populations 48 hours post 4-OHT removal. **C)** Flow cytometry analysis of dissociated day 16 human cultures immunostained for MNX1 and ISL1/2 shows that the vast majority of ISL1/2-positive cells are MNX1-positive and vice versa in both RFP^+^ and RFP^-^ populations. **D)** Day 18 human cultures immunostained for NEUN and ISL1/2 shows that the vast majority of NEUN-positive cells are ISL1/2 positive. (Scale bars = 50um)

**Supplemental figure 4:**
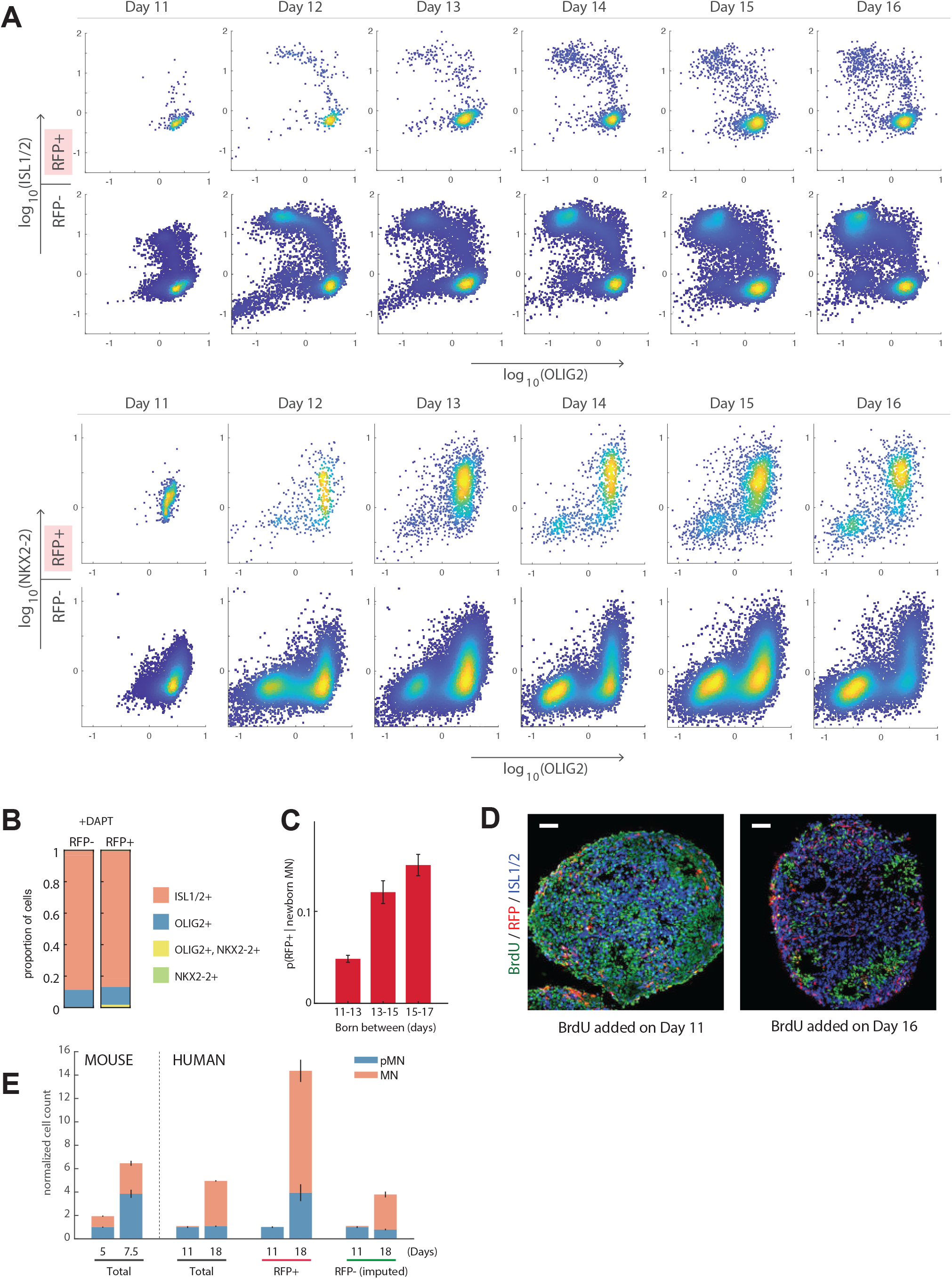
**A**) Flow cytometry plots for days 11-16 human cultures immunostained for ISL1/2, OLIG2 and NKX2-2 shows progressive increase in ISL1/2 expression in both RFP^+^ and RFP^-^ populations. **B)** Proportions of ISL1/2^+^, OLIG2^+^, OLIG2^+^/NKX2-2^+^ and NKX2-2^+^ cell types in day 16 cultures following DAPT treatment on day 11 shows that ~90% of both RFP^+^ and RFP^-^ populations are ISL1/2-positive. **C)** Proportion of RFP+ cells within newborn motor neuron population (as determined by BrdU labeling) increases over time. **D)** Day 18 human cultures with BrdU added on day 11 (left) or day 16 (right), immunolabeled for BrdU and ISL1/2. (Scale bar s= 50um) **E)** Relative numbers of motor neuron progenitors and motor neurons at the onset of and tail-end of neurogenesis in both mouse and human (normalized to number of progenitor cells at the onset of neurogenesis).

**Supplemental figure 5:**
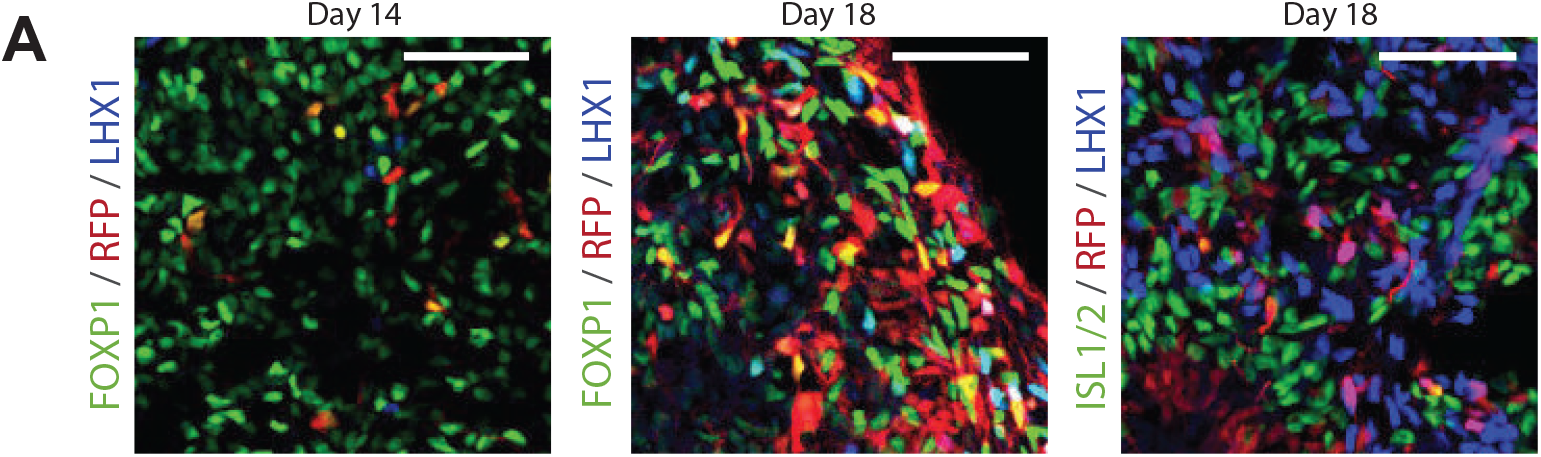
**A)** Left: Day 14 human cultures immunostained for FOXP1 and LHX1 shows that LHX1 is not yet expressed, but by day 18 is robustly expressed and overlaps with FOXP1, indicating that these cells are likely LMCl subtypes (middle). Right: LHX1-positive cells have lower ISL1/2 expression, further indicating that they are LMCl motor neurons. (Scale bars = 50um)

## Notes

### Competing Interest Statement

The authors have declared no competing interest.

